# Homology modeling and functional characterization of multidrug effluxor Mta protein from *Bacillus Atrophaeus*: An explanatory insilico approach

**DOI:** 10.1101/2020.12.29.424731

**Authors:** Mohammad Rejaur Rahman, Ishtiak Malique Chowdhury, Anik Banik, Emran Hossain Sajib

## Abstract

Phenotypically similar to *B. subtilis, Bacillus atrophaeus* is a Gram-positive, aerobic, spore-forming bacteria. It is a black-pigmented bacterial genus. Therefore, it is of interest to study the uncharacterized proteins in the genome. For a detailed computational sequence-structure-function analysis using available data and resources, an uncharacterized protein Mta (AKL87074.1) in the genome was selected. In this study, attempts were made to study the physicochemical properties, predict secondary structure, modeling the 3-D protein, pocket identification, protein-protein interaction and phylogenetic analysis of Mta protein. The predicted active site using CASTp is analyzed for understanding their multidrug resistance function. Because Mta is a MerR family member, these investigations on these functional aspects could lead us for better understanding of antibiotic resistance phenomenon.

## Introduction

The development and use of antibiotics has been one of the most important steps towards controlling infectious diseases in the 20th century. However, the subsequent appearance and spread of antibiotic resistance in pathogenic organisms have made many currently available antibiotics ineffective (Moellering Jr, 1998). So, Bacterial multidrug resistance (MDR) is a growing threat to human and animal health. To successfully fight the increasing numbers of drug resistant and multidrug-resistant (MDR) bacteria, extensive knowledge of the molecular mechanisms underlying microbial antibiotic resistance is required.

Microorganisms have developed various ways to resist the toxic effects of antibiotics and other drugs (Neu, 1992). One of these mechanisms involves the efflux of structurally and chemically diverse compounds, including antibiotics, antiseptics, and disinfectants, by membrane-bound multidrug transporters (Van Veen et al., 1999). The natural function of these proteins lies in protecting bacteria from external toxic chemicals. Indeed, these transporters have been found to provide enteric bacteria with natural resistance to bile salts (Baranova et al., 1999). On the basis of bioenergetic and structural criteria, multidrug transporters can be divided into two major classes. Secondary multidrug transporters utilize the transmembrane electrochemical gradient of protons or sodium ions to drive the extrusion of drugs from the cell. ATP-binding cassette (ABC)-type multidrug transporters use the free energy of ATP hydrolysis to pump drugs out of the cell.

Multiple multidrug transporters have been identified in Bacillus subtilis (Ohki and Murata, 1997; Kim et al., 2009), Escherichia coli (Edgar and Bibi, 1997), Pseudomonas aeruginosa (Kohler et al., 1997), Lactococcus lactis (Lubelski et al., 2006; van Veen et al., 1996), *Bacillus atrophaeus* UCMB-5137 (Baranova et al., 1999) Staphylococcus aureus (Floyd et al., 2010), Streptococcus pneumonia (Hashimoto et al., 2013; Boncoeur et al., 2012), Mycobacterium fortuitum (Aínsa et al., 1998), Mycobacterium tuberculosis (Aínsa et al., 1998) and Corynebacterium glutamicum (Yang et al., 2014). In *Bacillus subtilis*, there are two multidrug transporters namely Bmr and Blt, which belong to secondary multidrug transporters class. Mta (multidrug transporter activation), a member of the MerR family of bacterial regulatory proteins, is a global activator of *B. subtilis* multidrug transporter genes and constitutively activates transcription of Bmr and Blt, another putative membrane protein gene (ydfK) and its own gene (Godsey et al., 2001). MerR family members are dimeric proteins (Newberry & Brennan, 2004) and the majority of regulators in this family are activated in response to stress signals in bacteria, such as oxidative stress, heavy metals, cytotoxic compounds or antibiotics (Brown et al., 2003). So, activation of Mta ptotein activated drug transporters such as Bmr and Blt which successfully efflux the drug compounds out of the cell and exhibit resistance to drugs and antibiotics. To fight against the antibiotic resistance and drug resistance challenges, MerR family of regulatory protein like Mta protein is now a research interest.

In silico analysis of genes and proteins has been receiving greater attention with particular emphasis to find suitable biomarkers for rapid identification of different pathogenic genera (Kumar et al., 2016), designing of drugs to combat the pathogenic microbes and superbugs (Karumuri et al., 2015), diagnosis of infectious diseases (Kalia & Kumar, 2015), structural and functional analysis of proteins and enzymes (Pramanik et al., 2017), and discovery of potent microbial representative useful for several agricultural and animal feed industries (Verma et al., 2016; Pramanik et al., 2017). In the present study Mta protein of *Bacillus atrophaeus* UCMB-5137 were used for in silico analysis. *Bacillus atrophaeus* UCMB-5137 is an aerobic, grampositive, endospore-forming, rod-shaped bacterium whose description is nearly just like that of *Bacillus subtilis* except for the production of a pigment on media containing an organic source of nitrogen. (Nakamura, L. K. 1989). Attempts were made to study the physicochemical properties, predict secondary structure, modeling the 3-D protein, pocket identification, protein-protein interaction and phylogenetic analysis of Mta protein.

## Materials and Methods

### Recovery of the Targeted Sequence

The NCBI database (http://www.ncbi.nlm.nih.gov/) gives access to Biomedical and genomic data of numerous living beings. This database was utilized for the recovery of FASTA sequence of Mta protein. The FASTA format of the sequence was used for further in silico analysis.

### Analysis of Physico-Chemical Properties

Physico-chemical properties of the targeted protein sequence was analyzed by the ProtParam (http://web.expasy.org/protparam/) (Gasteiger, 2005) tool of ExPASy server. Utilizing this tool we analyzed molecular weight, theoretical pI, aliphatic index (AI), instability index, extinction co-efficients, GRAVY (grand average of hydropathy), amino acid composition, atomic composition etc.

### Secondary Structure Prediction

The secondary structures combine to form the tertiary (3D) structure that determines the function of a protein. PSIPRED v3.3 (http://bioinf.cs.ucl.ac.uk/psipred/) (Liam J. McGuffin, 2000) and CFSSP: Chou and Fasman Secondary Structure Prediction server (http://cho-fas.sourceforge.net/) (Kumar, 2003) were used to predict the overall secondary structure of Mta protein and to count the number of helices, sheets and turns.

### Homology Modeling, Refinement and Model Quality Assessment

The I-TASER server (Yang et al., 2015) was utilized for determining the 3D structure of Mta protein based on the degree of similarity between the target protein and available template structure from PDB. Refinement was performed using GalaxyWEB server (http://galaxy.seoklab.org/) (Ko et al., 2012). The refined protein structure was validated through the Ramachandran plot assessment by the RAMPAGE software (http://mordred.bioc.cam.ac.uk/~rapper/rampage.php) (Lovell et al., 2003) and the quality factor was assessed by using ERRAT server (https://servicesn.mbi.ucla.edu/ERRAT/) (Colovos & Yeates, 1993).

### Sub-cellular Localization and active site determination

Sub-cellular localization determination is crucial for understanding protein function and is also vital for genome analysis. Sub-cellular localization of the Mta was determined by CELLO version 2.5 (Yu et al., 2004) which is a multiclass support vector machine classification system. Another two servers ngLOC (King & Guda, 2007) and PSIpred (Buchan & Jones, 2019) were also used for confined conformation. The generated 3D model was further used for finding pocket to identify residues which are involved in binding of substrate or transcription factor. Active site of the protein was determined by the computed atlas of surface topography of proteins (CASTp) server (Tian, 2018).

### Functional Analysis

The Mta protein was analyzed for the presence of conserved domains based on sequence similarity search with near orthologous family individuals. For this purpose, NCBI database (http://www.ncbi.nlm.nih.gov/cdd/) was used (Marchler-Bauer, 2014). To identify the super family and family of the Mta protein, SUPERFAMILY 2.0 server (http://supfam.org/) was used (Gough, 2001). In order to find the motif present in Mta protein Motif Search tool (https://www.genome.jp/tools/motif/) was used (Kanehisa, 2002). Protein–protein interaction (PPI) plays an amazingly momentous part within the development, propagation, and digestion system of all lives. To know the interaction of Mta protein of *Bacillus atrophaeus* UCMB-5137 with other closely related proteins STRING v11 (http://string-db.org/) server (Szklarczyk, 2018) was used.

### Phylogenetic Analysis

Phylogenetic analysis is vital since it improves our understanding of how genes, genomes, species (and molecular sequences more generally) evolve. The UPGMA tree was constructed for focused on protein utilizing MEGA software (Kumar, 2018).

## Result

### Sequence retrieval

An Mta protein from the species *Bacillus atrophaeus* UCMB-5137 with accession number AKL87074.1 has been retrieved from NCBI database that comprise of 244 amino acids for this computational study.

### Analysis of physico-chemical properties

Physiochemical characterization is very crucial to characterize a specific protein. Table 1 shows the different physiochemical properties of Mta protein of *Bacillus atrophaeus* UCMB-5137. Isoelectric point (pI) is the pH at which the surface of protein is covered with charge but net charge of protein is zero. Isoelectric point (pI) of Mta protein is 5.44. So it seemed acidic protein. Average molecular weight of the protein is 28961.95 Da. The instability index (II) of the protein is 44.04 which classifies this protein may be unstable in nature (), but on contrary the protein showed higher aliphatic index (80.70) which suggested that the protein is thermostable. The GRAVY value for this protein is very low (−0.697) and Lower value of GRAVY shows that the proteins have better interaction with water.

**Table 1:**
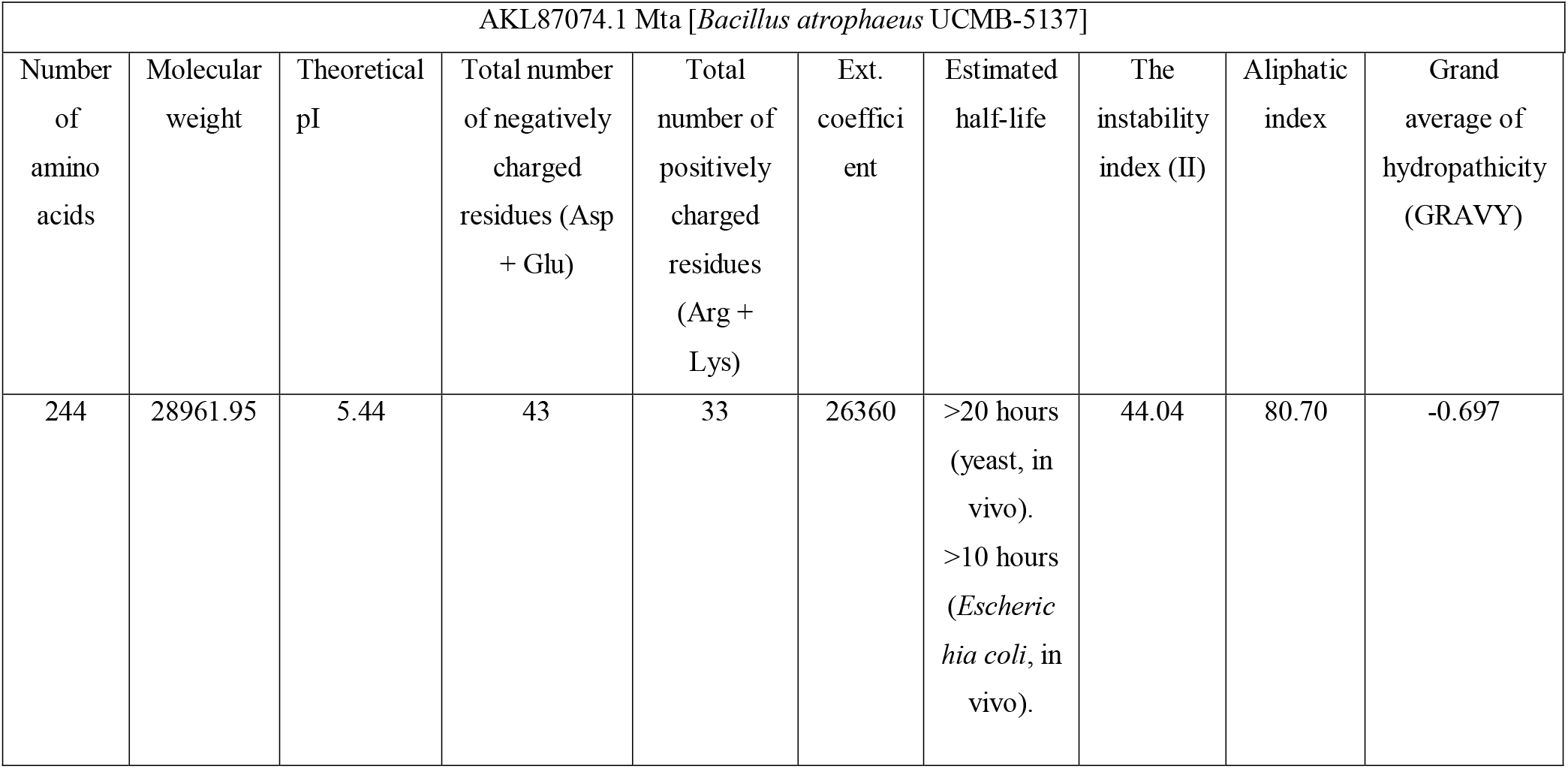
Physiochemical analysis.

**Table 2:** Subcellular Localization of Mta.

| CELLO | ngLOC | PSIpred |
| --- | --- | --- |
| Cytoplasmic | Cytoplasmic | Cytoplasmic |

### Secondary Structure Prediction

The secondary structure is the intuitive that happen between the C, O, and NH bunches on amino acids in a polypeptide chain to create α-helices, β-sheets, turns, circles, and other shapes. The percentile contribution of three classes of secondary arrangements viz. Helices, Sheets and Turns are 84%, 72.5% and 12.7% respectively which is deduced from the web server (Fig. 1). This result indicates that helices>sheets>turns in the protein. Disordered protein binding site was not detected (Fig.1). Secondary arrangements indicates that the Mta protein is folded, which indicates the stable nature of protein. Here, a very higher percentage of α-helical conformation indicates that the protein is thermostable (Pramanik et al. 2017). The limitations of X-ray Crystallography and NMR are overcome by the prediction of secondary structure elements of protein (Roy et al., 2015). In silico characterization including secondary structure prediction of ACC deaminase protein of Mesorhizobium were done by Pramanik et al. (2017), the phytase of different Bacillus spp. by Verma et al. (2016). Roy et al. (2015) reported that the prediction of secondary structural elements was vital to detect the conformational changes within the protein of interest.

**Figure 1.**
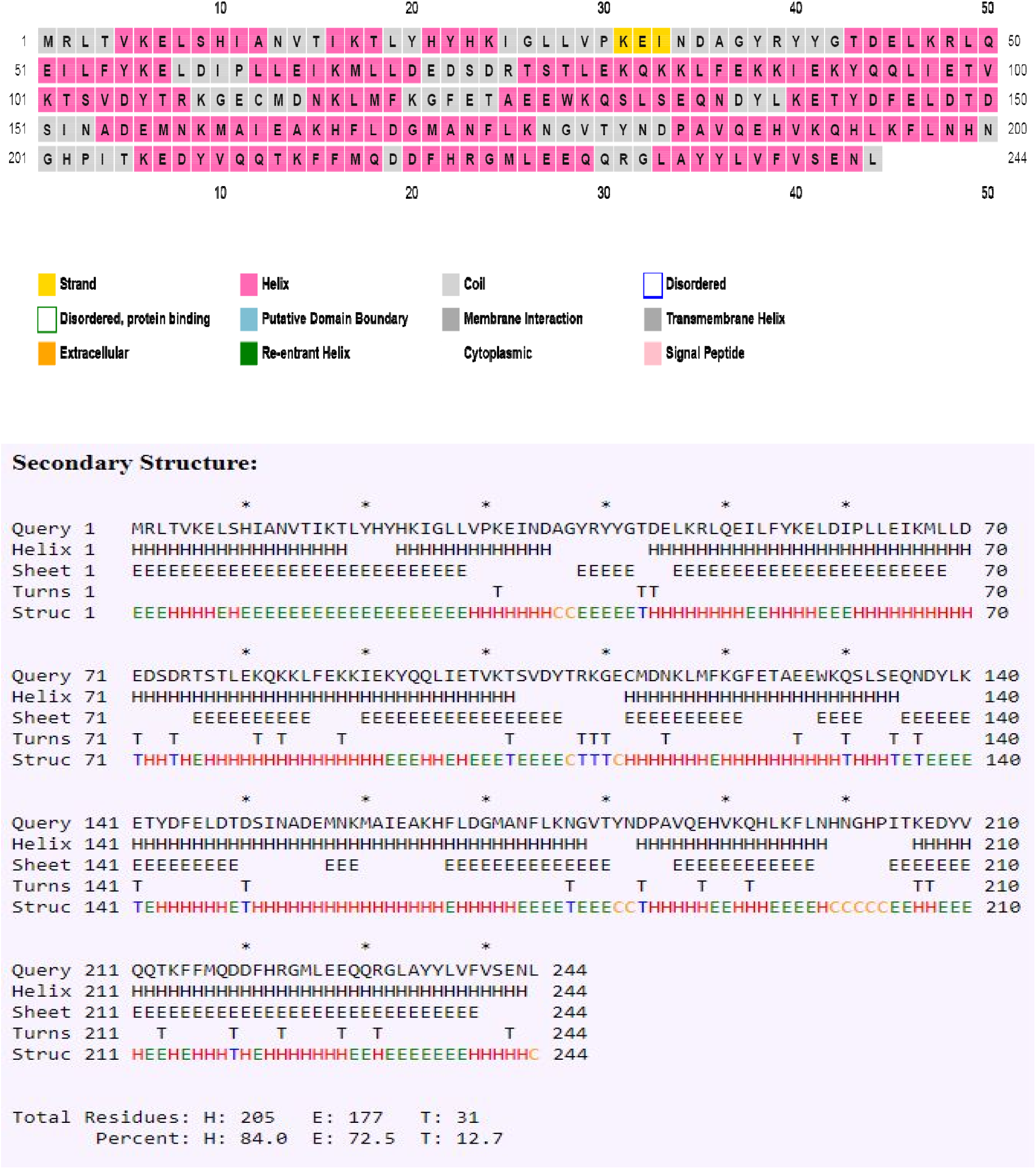
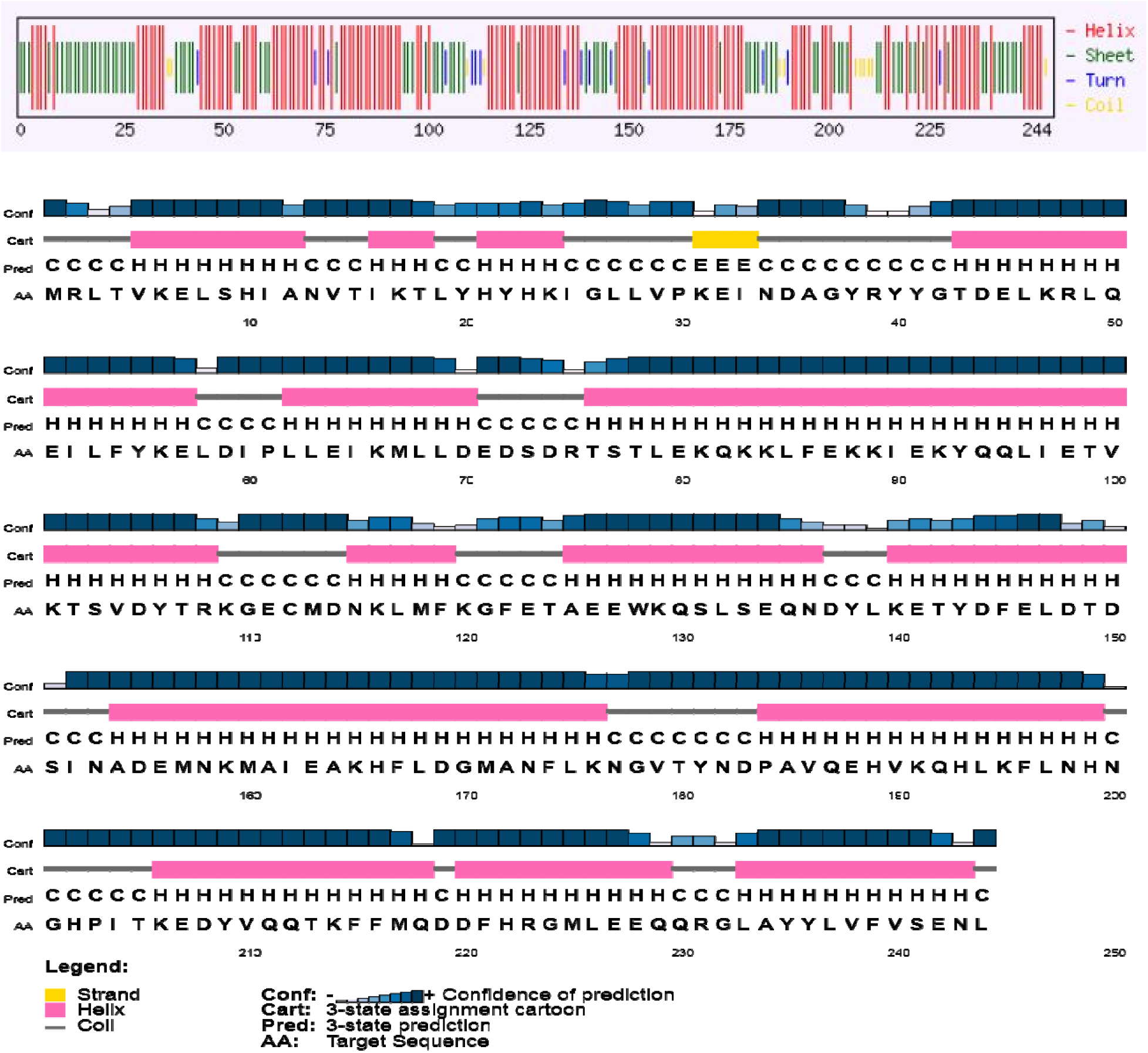
Secondary structure analysis of Mta protein.

### Homology Modeling

I-TASSER predicted five models for Mta protein of *Bacillus atrophaeus* UCMB-5137, which were ranked based on cluster size. The LOMETS threading programs selected ten best templates (with the highest Z-score) that were used to predict the tertiary structures. Homology modeling was performed by using 3qaoA from RCSB Protein Data Bank as a best suited template for Mta protein. Results showed that model 1 had the highest C-Score of −3.01 while the estimated TM-score and RMSD were 0.37±0.13 and 12.9±4.2Å (Fig. 2). After refinement by RAMPAGE, 80.6% and 14.5% residues were in the favored and allowed region revealed by Ramachandran plot analysis (Fig. 2). The ERRAT server showed that this predicted model has a quality factor of 62.66. In silico based homology modeling was also shown by several workers [17–19, 31, 33] to predict a variety of 3D protein models of interest.

**Figure 2.**
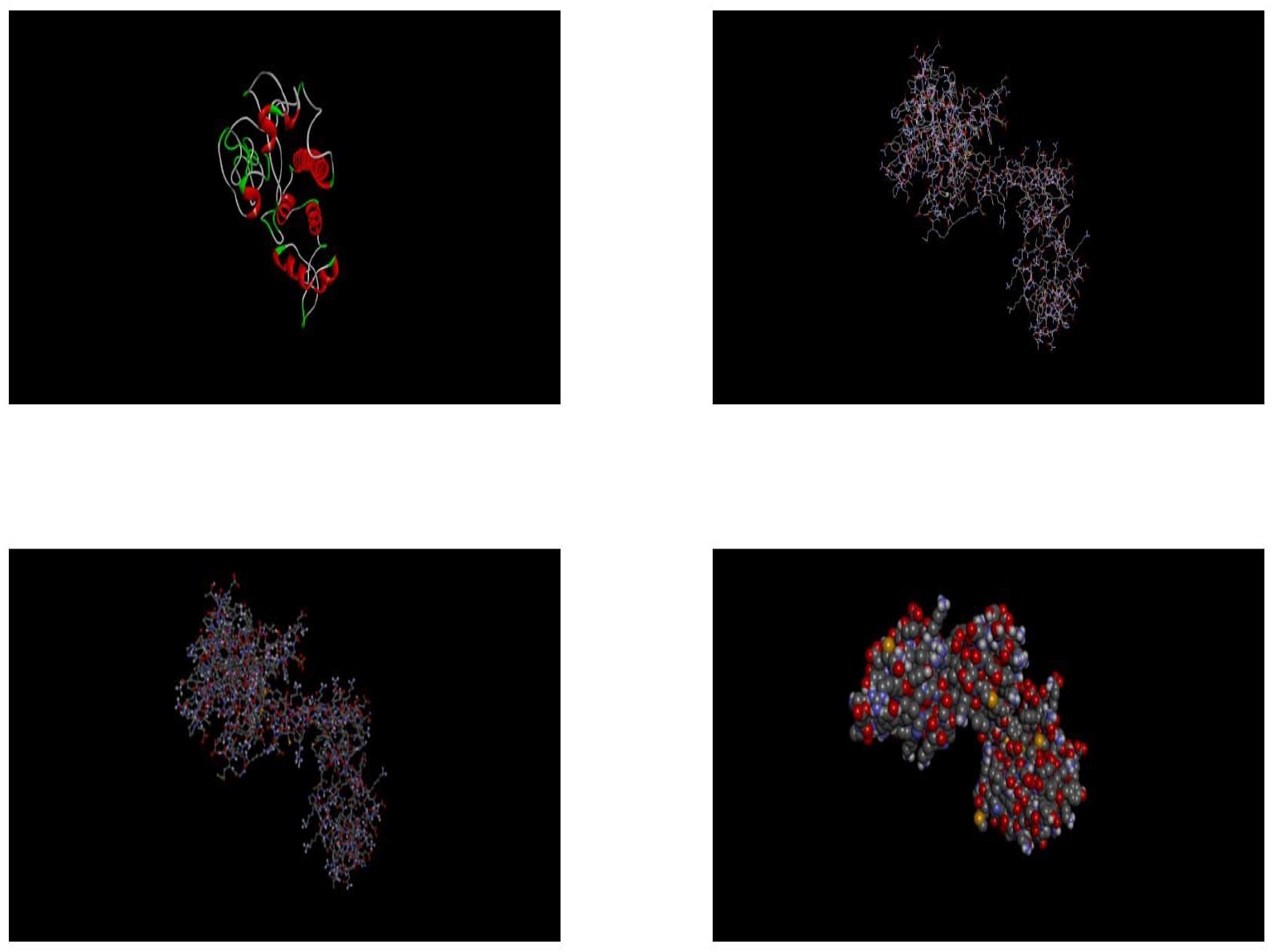
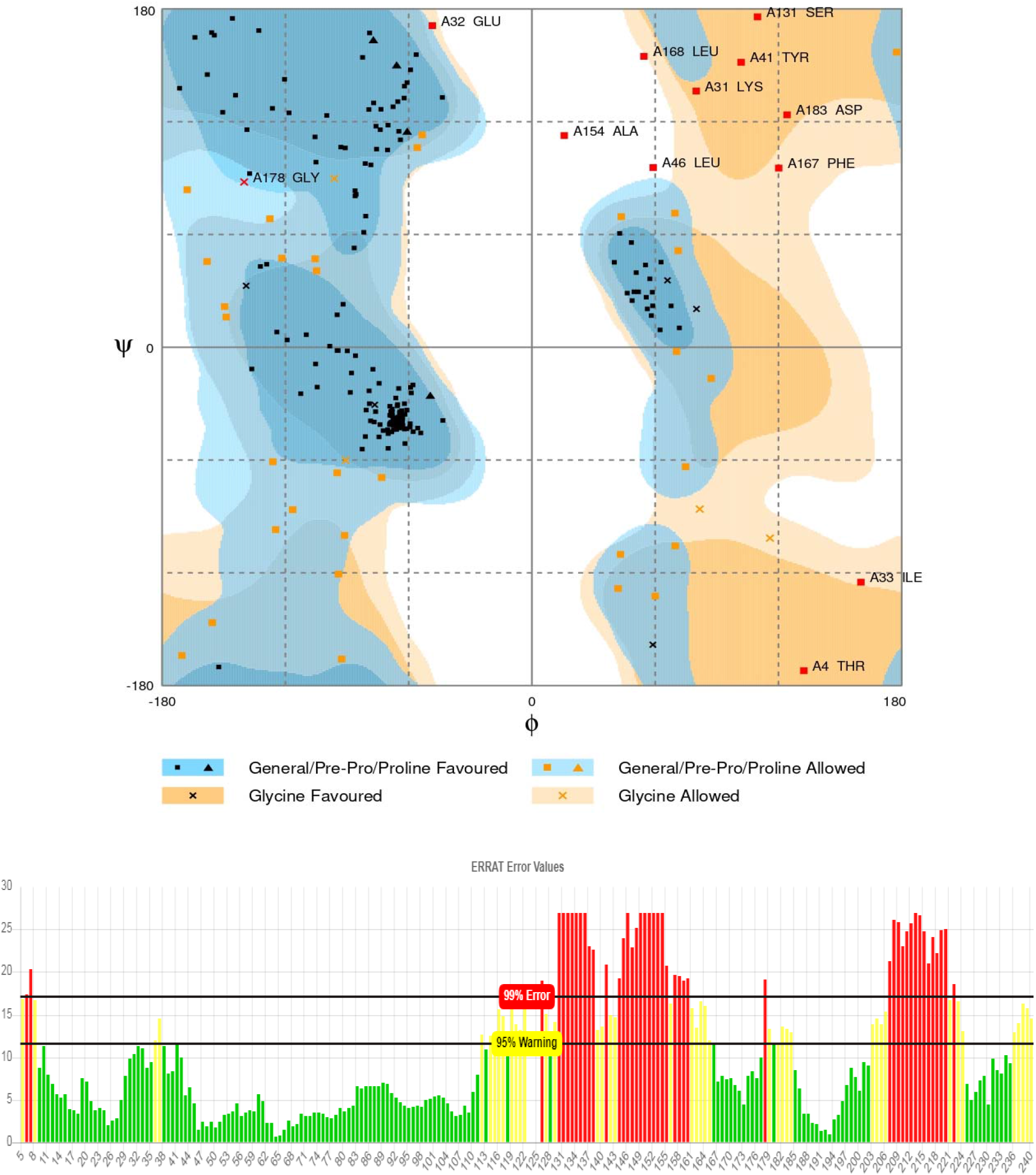
3D structure, Ramachandhan plot, ERRAT value.

### Sub-cellular localization

Sub-cellular localization is an essential feature of a protein. Cellular functions are usually localized in specific enclosed area; so, foretelling the sub-cellular localization of an unknown protein may possibly use to obtain handy information about their function. Therefore, this information is also valuable for drug designing for the target protein (Wang et al., 2005) For determining the sub-cellular localization o Mta protein we used three online based server and all of these three servers suggested that the location of Mta protein is in the cytoplasm.

The identification and characterization of functional sites on proteins have increasingly become an area of interest. On account of the analysis of the active site residues for the binding of ligands provides insight towards the design of inhibitors for the protein. Result of CASTp sever showed that there are 26 active sites (supplementary file 1) of this protein. The volume and area of the best pocket which identified by using CASTp server is volume (SA) is 1553.115 and area (SA) is 374.141.

**Figure 3.**
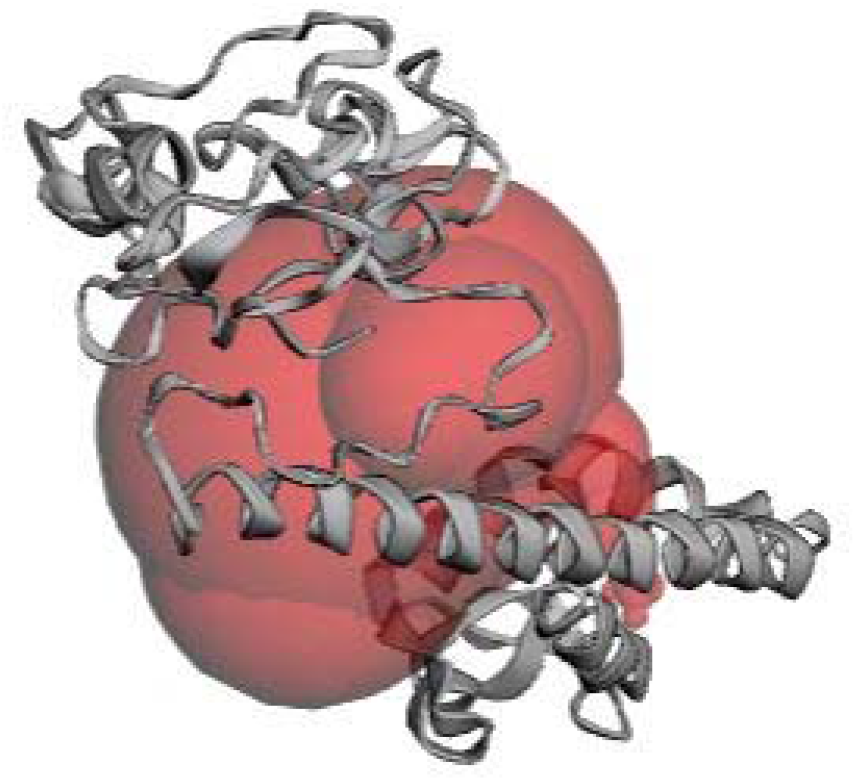
Active site determined by CASTp.

### Functional Analysis

The Conserved Domain Database of NCBI is a resource for the annotation of functional units in proteins which revealed that Mta protein has two domains. These are Helix-Turn-Helix DNA binding domain (HTH_TipAL-Mta) and TipAS antibiotic-recognition domain (TipAS) (Fig. 4). This TipAS domain is found at the C-terminus of some MerR family protein. Based on a collection of hidden Markov models, the SUPERFAMILY 2.0 server showed that Mta protein has two superfamily and two families (Fig. 5). These families are DNA-binding N-terminal domain of transcription activators and Antibiotic binding domain of TipA-like multidrug resistance regulators. Biological sequence motifs are characterized as brief, as a rule, settled length, sequence designs that will speak to vital structural or useful highlights in nucleic acid and protein sequences such as transcription binding sites, splice junctions, active sites, or interaction interfaces. Result of Motif Search tool showed that Mta protein of *Bacillus atrophaeus* UCMB-5137 has five motifs (Fig. 6) (supplementary file 2). STRING visualizes weighted and integrated and a confidence score of protein’s functional associations in a network of genome-wide connectivity (Szklarczyk, 2016). The STRING server has detected three interacting proteins viz., ADP34599.1, DnaJ and ADP31438.1, where ADP34599.1 and ADP31438.1 are Transcriptional regulators (Fig. 7). Among them DnaJ protein is a chaperone protein which participates actively within the reaction to hyperosmotic and heat shock through preventing the aggregation of stress-denatured proteins and by disaggregating proteins.

**Figure 4:**
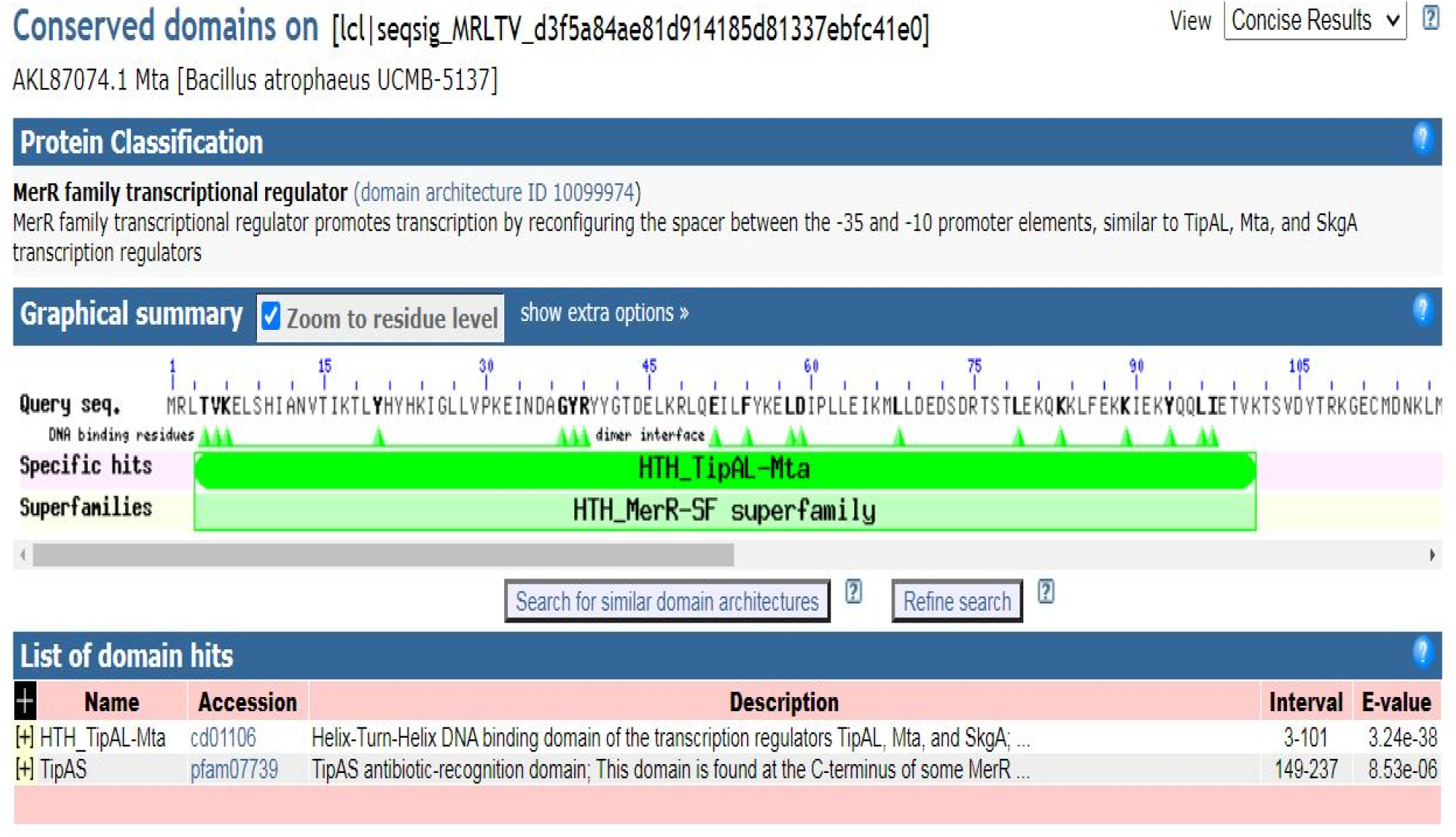
Domain search using NCBI CDD.

**Figure 5:**
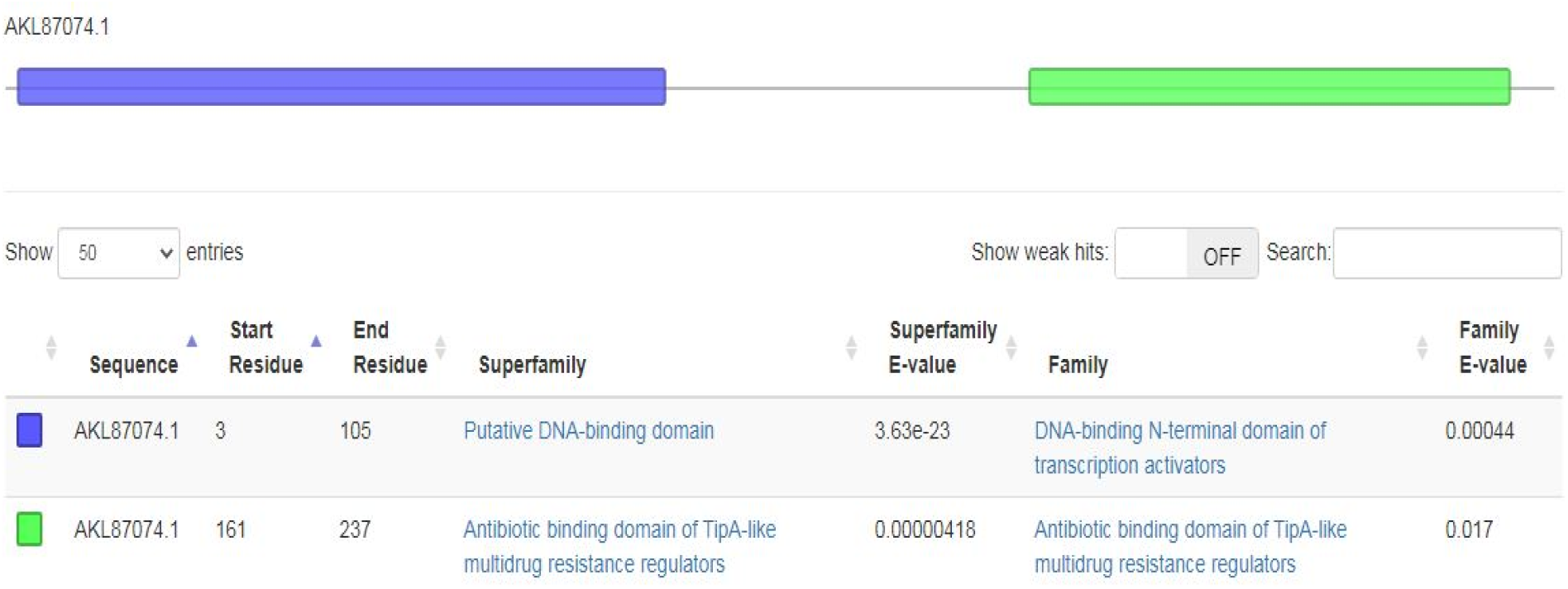
Superfamily and Family of Mta protein.

**Figure 6:**
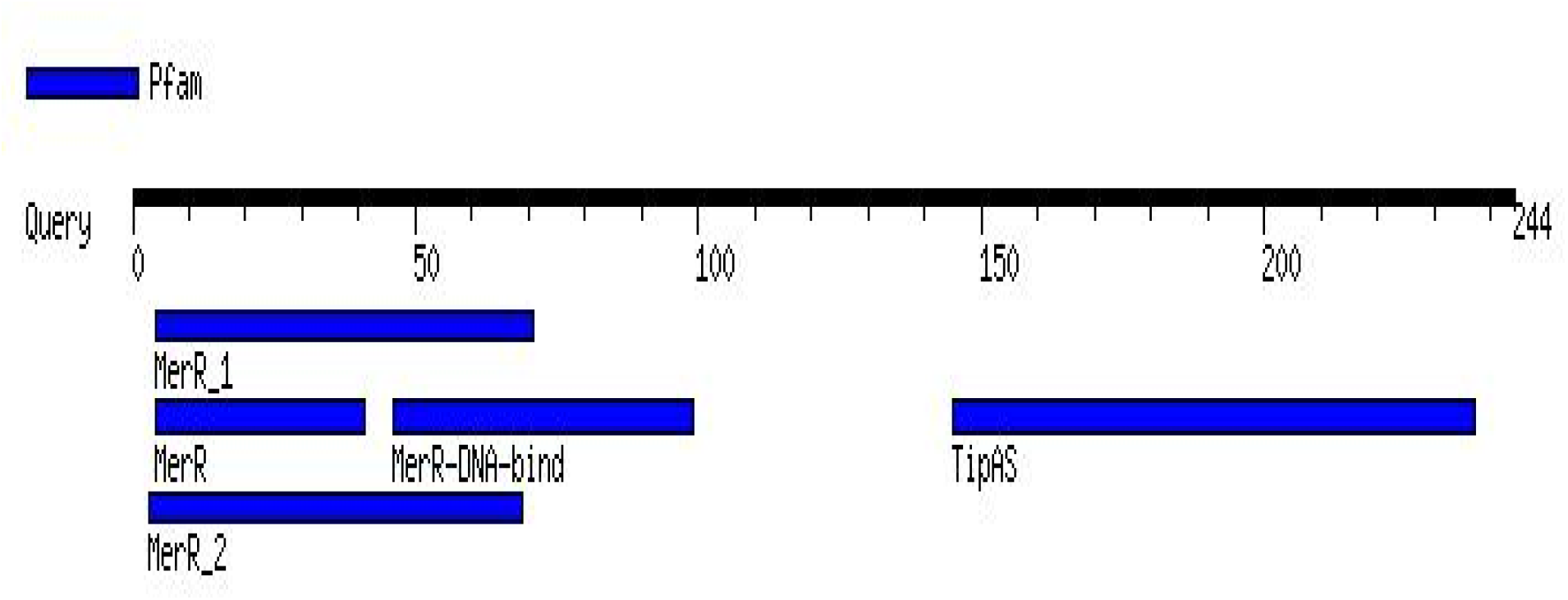
Motifs found using motif finder.

**Figure 7.**
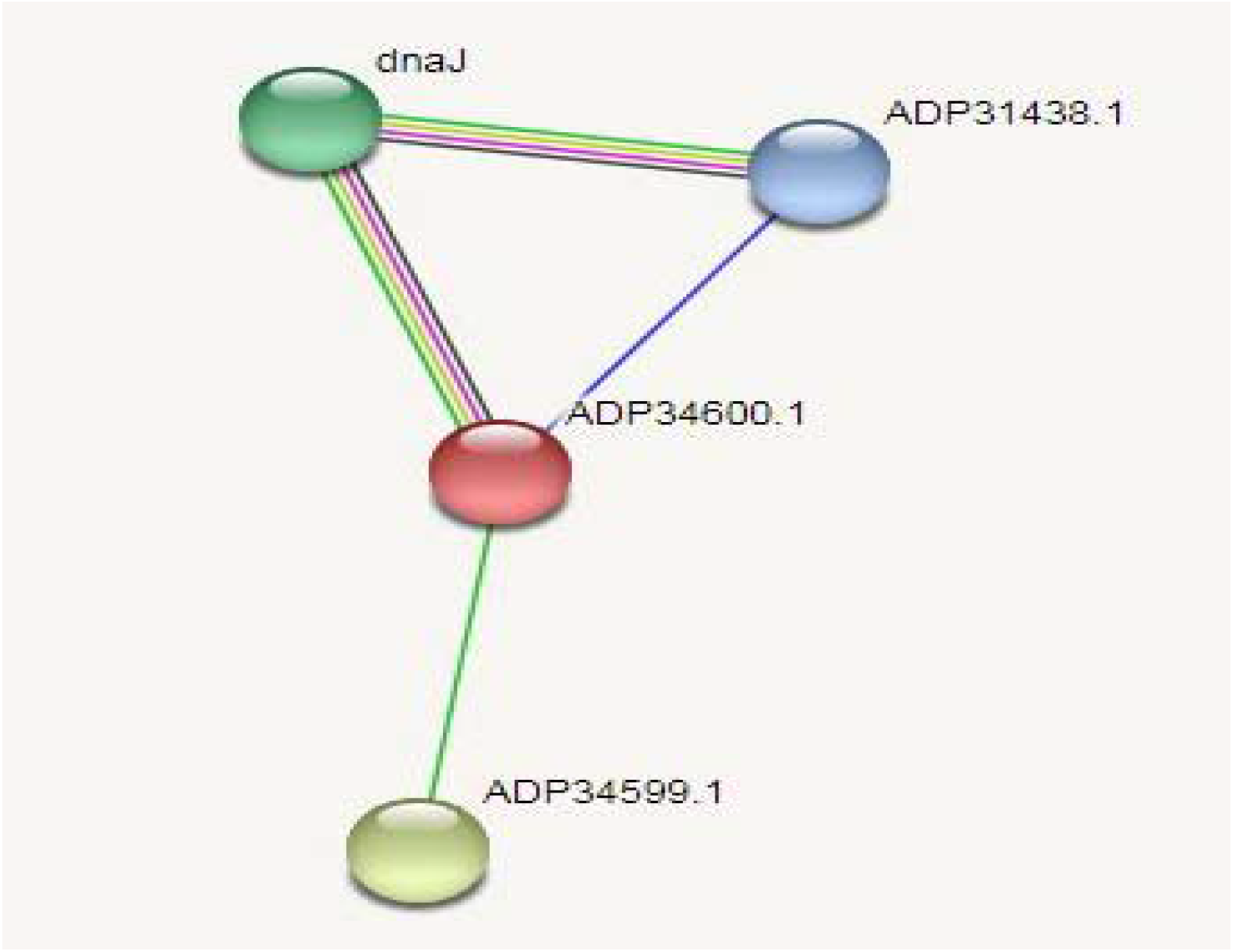
Protein-protein Interaction analysis using STRING.

### Phylogenetic Analysis

Phylogenetics is vital since it improves our understanding of how genes, genomes, species (and molecular sequences more generally) evolve. The UPGMA tree was contructed for focused on protein utilizing MEGA software (The optimal tree with the sum of branch length = 0.22278011 is shown. The tree is drawn to scale, with branch lengths in the same units as those of the evolutionary distances used to infer the phylogenetic tree. The evolutionary distances were computed using the Poisson correction method and are in the units of the number of amino acid substitutions per site. This analysis involved 15 amino acid sequences. All ambiguous positions were removed for each sequence pair (pairwise deletion option). There were a total of 245 positions in the final dataset.

**Figure 8:**
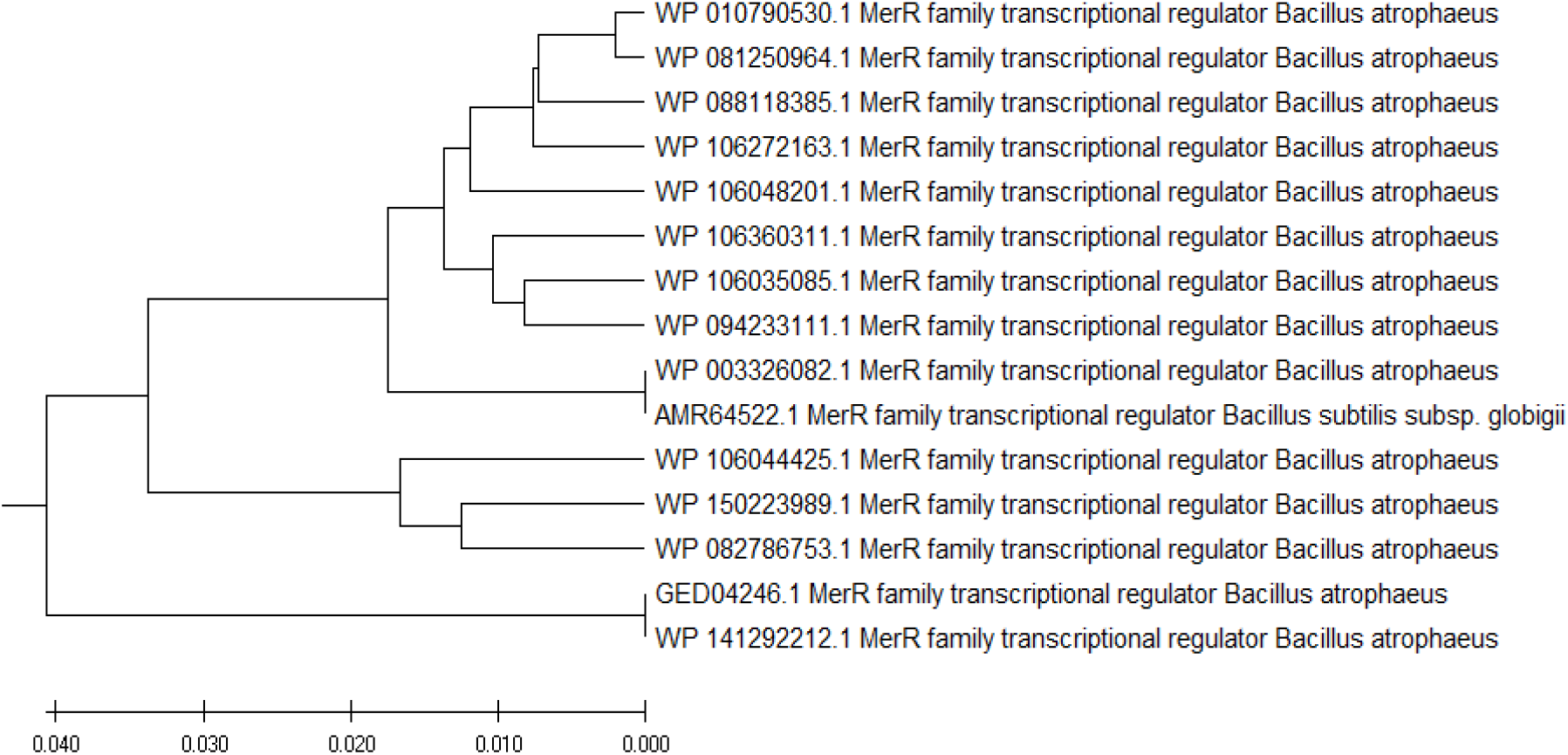
Phylogenetic Analysis using MEGA.

The present work might be shown some important aspects in future direction in computational studies. Firstly, this study demonstrates about the structural annotation of Mta protein which is now-a-days getting evoking interest to the present reaserachers because of its function in antibiotic resistance mechanism (Musarrat et al., 2009). Secondly, knowing the nature of the protein such as cytosplasmic, thermostable and folded might help to work out wet lab experiments (Li et al., 2015) less effortlessly limiting the number of frequent trials and errors etc. Thirdly, the active site of Mta protein and protein interaction analysis gives us an overview on the function of Mta protein. This in silico based information about Mta protein will play an important role in further researcher related antibiotic resistance mechanism. Through utilization of this data researcher can propose some supportive drug during antibiotic treatment by docking analysis to improve efficacy of antibiotic by prohibiting the activation of Mta protein. Activation of Mta protein is responsible for activating multidrug efflux proteins inside the cell.

## Conclusion

Mta protein, one of the key proteins present in different microorganisms, plays important role in antibiotic resistance mechanism. Therefore, understanding the structural properties of this protein is important to get further insight into the functional aspect and interaction with other proteins and molecules. From the in silico characterization of this study, it was revealed that Mta protein is a thermostable, acidic, and a molecular mass of 28961.95 Da having mainly two conserved domains. This work might be a valuable contribution in the field of Bioinformatics research and may help other researchers to get an idea about the protein structure, its physicochemical properties, structural motifs, and protein protein interaction essential for pharmaceutical and health sector.

---

**Supplementary Table 2:** Motifs with Independent E-value.

| **Pfam** | **Independent E-value** | **Description** |
| --- | --- | --- |
| <u>MerR_1</u> | 4..71(3e-12) | PF13411, MerR HTH family regulatory protein |
| <u>MerR</u> | 4..41(3.3e-11) | PF00376, MerR family regulatory protein |
| <u>TipAS</u> | 145..237(1.5e-10) | PF07739, TipAS antibiotic-recognition domain |
| <u>MerR-DNA-bind</u> | 46..99(0.0002) | PF09278, MerR, DNA binding |
| <u>MerR_2</u> | 3..69(0.1) | PF13591, MerR HTH family regulatory protein |

**Supplementary Table 2:** Pockets with area and volume.

| Pocket ID | Area (SA) | Volume (SA) |
| --- | --- | --- |
| 1 | 374.141 | 1553.115 |
| 2 | 494.368 | 407.104 |
| 3 | 169.632 | 73.266 |
| 4 | 80.306 | 20.749 |
| 5 | 21.781 | 6.768 |
| 6 | 28.810 | 6.515 |
| 7 | 11.698 | 4.145 |
| 8 | 14.951 | 3.757 |
| 9 | 13.712 | 2.221 |
| 10 | 11.145 | 1.755 |
| 11 | 6.359 | 1.436 |
| 12 | 7.012 | 1.397 |
| 13 | 3.082 | 0.848 |
| 14 | 5.136 | 0.562 |
| 15 | 1.781 | 0.439 |
| 16 | 1.709 | 0.072 |
| 17 | 1.541 | 0.071 |
| 18 | 0.816 | 0.039 |
| 19 | 0.624 | 0.022 |
| 20 | 0.764 | 0.019 |
| 21 | 0.744 | 0.011 |
| 22 | 0.336 | 0.007 |
| 23 | 0.251 | 0.007 |
| 24 | 0.385 | 0.007 |
| 25 | 0.018 | 0.000 |
| 26 | 0.002 | 0.000 |

